# Selection on sporulation strategies in a metapopulation can lead to coexistence

**DOI:** 10.1101/2024.02.17.580810

**Authors:** Stephen R. Proulx, Taom Sakal, Kelly Thomasson

**Affiliations:** Department of Ecology, Evolution, and Marine Biology, UC Santa Barbara, Santa Barbara, CA 93106, USA

**Keywords:** metapopulation, extinction, sporulation

## Abstract

In constant environments the coexistence of similar species or genotypes is generally limited. In a metapopulation context, however, types that utilize the same resource but are distributed along a competition-coexistence trade-off, can coexist. Much thought in this area focuses on a generic trade-off between within-deme competitive ability and between-deme dispersal ability. We point out that the sporulation program in yeasts and other microbes can create a natural trade-off such that strains which initiate sporulation at higher rates suffer in terms of within-deme competition but benefit in terms of between deme dispersal. We develop simple metapopulation models where the within-deme behavior follows simple chemostat dynamics. We first show that the rate of sporulation determines the colonization ability of the strain, with colonization ability increasing with sporulation rate up to a point. Metapopulation stability of a single strain exists in a defined range of sporulation rates. We then use pairwise invasability plots to show that coexistence of strains with different sporulation rates generally occurs, but that the set of sporulation rates that can potentially coexist is smaller than the set that allows for stable metapopulations. We then extend our pairwise results to show how a continuous set of strains can coexist and verify our conclusions with numerical calculations. Our results show that stable variation in sporulation rates is expected under a wide range of ecological conditions.

## Introduction

Ecology has long explored the processes that can produce diverse communities of coexisting species (Hutchinson, 1959; Brown, 1981). When species utilize different aspects of the environment, either by consuming different resources (Ackermann and Doebeli, 2004; Leimar et al., 2013), avoiding predators (Roughgarden and Feldman, 1975), or through any other mechanism that increases niche differences (Letten et al., 2018; Chesson, 2000). Empirically, there is often more coexistence of similar species than can be explained by modern coexistence theory (MCT) (Ellner et al., 2019), and in some cases this is revealed to be due to hidden physiological differences (Burson et al., 2019).

Even when species compete over a single resource, they may coexist through trade-offs that partition space and time, such as when the competition colonization trade-off holds (Levins, 1966; Hastings, 1980; Tilman, 1994). We examine how changes in the sporulation strategy of yeast cells can create a competition-colonization trade-off and allow coexistence of yeast strains (or species) that are otherwise physiologically identical. This opens a new avenue for explaining coexistence of similar species.

The competition-colonization trade-off was promoted by Tilman (1994) based on the Levins-Hastings competitor-prey model (Levins, 1966; Hastings, 1980). In Tilman’s formulation, a ranked set of competitors is considered, where species with higher within group competitive ability have lower dispersal ability. The Tilman model assumed that competition within the patch was deterministic and had time to play out before additional colonization events could occur, which leads to a strict replacement of lesser competitors within local demes. Two competitors can coexist when the lesser competitor has a sufficiently greater colonization ability, and the inclusion of a lesser competitor never leads to the extinction of the better competitor (Tilman, 1994). This strict competitive ranking leads to some unusual features of the model, in that the most competitive strain can always invade but reaches an equilibrium metapopulation density that is virtually zero (Kinzig et al., 1999). This feature of the model has been explored and relaxed by allowing for competition functions that are smoother (Adler and Mosquera, 2000) and allowing for the existence of preemption effects (Yu and Wilson, 2001; Amarasekare, 2003; Calcagno et al., 2006). When competition is completely based on preemption, coexistence does not occur (Yu and Wilson, 2001). However, when there is a partial preemption effect, coexistence is generally favored but the set of coexisting types is affected (Calcagno et al., 2006).

As is elegantly described in Kinzig et al. (1999), the Hastings-Tilman model formulation allows the better competitor to create a competition “shadow” that prevents invasion of similar, but weaker, competitors. Better competitors, however, can always invade, and so we expect that successive introductions/mutations will lead to better and better competitors being part of the community. Kinzig et al. (1999) also explain that as the better competitor gets closer to the minimal colonization ability that allows metapopulation persistence, the competition shadow gets smaller, leading to dense packing of species along the competitive ability gradient. This leads to some paradoxical results, whereby the best competitor in the system has a patch occupancy frequency of approximately zero, but the total density of species within a window of competitive abilities follows a 3/2 power law, and the entire metapopulation is occupied. The extension of Calcagno et al. (2006) goes some way towards resolving this paradox by showing that there is a limit to the invasion of species that are close to this metapopulation persistence boundary. These methods have been extended to consider the relative importance of spatial heterogeneity and colonization trade-offs in a life-history framework (Amarasekare et al., 2004). Amarasekare et al. (2004) also call for the inclusion of mechanistic trade-off models and suggest that an adaptive dynamics approach may be helpful, which we take here.

In its natural habitat, the yeast *S. cerevisiae* primarily exist as vegetative mitotic diploid cells. They disperse to new environments through the digestive tracts of insect vectors (Gibbs and Stanton, 2001; Coluccio et al., 2008; Stefanini et al., 2012). Vegetative cells experience extreme mortality when ingested by insects, but spores do not (Coluccio et al., 2008). Yeast cells can go through a meiotic division process to produce haploid quiescent spores enclosed within a protective structure known as an ascus (Neiman, 2005). The initiation of sporulation occurs when diploid cells encounter unfavorable conditions, a process that can be triggered by changes in nutrient availability and pH, which are often correlated with resource consumption and waste production by yeast populations (Neiman, 2005). Sporulation rate known to depend on each strain’s evolutionary history, with more domesticated strains having lower rates of induced sporulation (Liti et al., 2009).

In our previous research, we investigated how budding yeast adapts to survive its passage through the digestive tracts of flies (Thomasson et al., 2021). We found that strict fly passaging led to the evolution of a sharpened transition to the spore state and an increase in spore production in each yeast strain. Notably, even domesticated yeast strains, initially less inclined to sporulate, exhibited substantial increases in spore production. Even though prior evolutionary history (i.e. domestication) constrained the evolved increase in sporulation, it was clear that mutations altering the sporulation program are readily accessible. We also found that the timing of sporulation was evolvable, showing that alternative types of sporulation plasticity are possible. Our focus in this work is to study the forces that can lead to evolution of the sporulation program in a metapopulation.

To motivate our model, imagine a vineyard where grapes occasionally fall to the ground and are split open. Each grape represents a patch, and sugar and nutrients seep out of the grape where yeast naturally grow. On any given grape the yeast strain with lowest sporulation rate is the strongest competitor and eventually drives all other strains to extinction. However, grapes are ephemeral things and disaster often befalls them. They may be removed by the farmer, a passing animal may eat them, or the may be washed away by rain. Any grape will eventually be destroyed, so it is not enough for a yeast strain to be a good competitor – it must be a good colonizer too. Only yeast genotypes that are able to be transferred to new grapes have a potential to maintain a presence in the metapopulation of crushed grapes.

Previous work has focused on the dispersal benefit of sporulation (Coluccio et al., 2008) and that sporulation before insect mediated dispersal could lead to higher rates of outcrossing creating an advantage in terms of adaptability of the yeast population (Reuter et al., 2007). The evolution of sporulation in a structured population has also been explored from a kin selection perspective (Ratcliff et al., 2013). In this framework, yeast who consume less resources by sporulating provide a benefit to their relatives that provides a net kin selection benefit. This has recently been expanded to consider the general kin selected effects of dormancy (Twyman and Gardner, 2023).

Our goal was to incorporate explicit eco-evolutionary dynamics into a metapopulation model of yeast sporulation and dispersal in order to study the evolution of the sporulation program. We derived a model of yeast population dynamics that accounts for the production of both vegetative cells and spores, and used the equilibrium density of spores to calculate the colonization ability of competing strains. We first take an adaptive dymamics approach to show how pairwise invasibility plots can be used to consider how sporulation evolves when the supply of mutations is low. We then consider a situation where many mutants are able to segregate at once and derive an integro-differential equation to describe the frequency distribution of sporulation mutants.

## Models

### Within patch dynamics

We model within-patch dynamics using a chemostat-like resource renewal model. In this scenario, resources are renewed at a constant rate and are removed (i.e. by natural degradation or outflow) at a constant per capita rate. Vegetative yeast cells consume the resource at a constant per-capita rate. Spores are produced at a constant rate at each cell division. While sporulation is known to respond to the level of resources in the media, we here model strategies that vary in their fixed rate of sporulation.

The dynamics can be described as

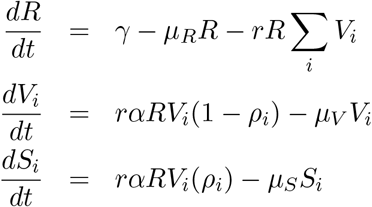

where *R* is the density of the resource, *γ* is the supply rate of the resource, *µ*_*R*_ is the natural degradation or removal rate of the resource, *V*_*i*_ is the density of vegetative cells of the *i*th genotype, *r* is the rate at which resource is consumed upon interaction between the yeast and the resource, *α* is the conversion efficiency of resource into yeast cells, *ρ*_*i*_ is the sporulation rate of the *i*th genotype, *µ*_*V*_ is the natural mortality rate of vegetative cells, *S*_*i*_ is the density of spores of the *i*th genotype, and *µ*_*S*_ is the natural mortality rate of spores (see table 1). The sporulation rate, *ρ* can also be thought of as the fraction of daughter cells that enter the sporulation pathway and become spores.

**Table 1:** Notation.

| Variable | Definition |
| --- | --- |
| **General parameters** |  |
| $\gamma$ | Supply rate of the resource |
| $\mu_R$ | Rate of removal of the resource |
| $R$ | Density of the resource |
| $r$ | Consumption rate per cell |
| $V_i$ | Density of vegetative cells of genotype $i$ |
| $\alpha$ | Conversion efficiency of resources into cells |
| $\rho_i$ | Sporulation rate of genotype $i$ |
| $\mu_V$ | Death rate of vegetative cells |
| $\mu_S$ | Death rate of spore cells |
| $P_i$ | Proportion of patches occupied by genotype $i$ |
| $c_i$ | Colonization rate of genotype $i$ |
| $\mu_P$ | The patch extinction rate |

This differs from a strict chemostat model in that the degradation rates of the different quantities are allowed to be independent (the *µ* parameters are not required to be equal for the different state variables). This assumes that the resource can leave the system by some degratory process that does not strictly involve flushing out of the system and may represent other biological or non-biological physical processes. The degradation rates of the vegetative cells and spores are assumed to be the combination of natural mortality (including for example senescence) and removal due to physical processes and depredation, which we do not explicitly model here.

Given a set of *n* genotypes within a patch, we index the genotypes such that *ρ*_1_ *< ρ*_2_ and *ρ*_*i*_ *< ρ*_*i*+1_. So long as 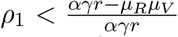 then genotype 1 persists in the population and all other genotypes go extinct (see appendix A). If *ρ*_1_ does not satisfy this criteria, then all yeast go extinct because the rate of vegetative cell death combined with spore production is too high.

The single stable equilibrium of this system has

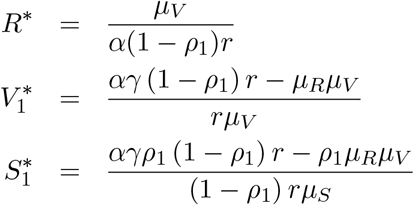

Generally, the equilibrium density of vegetative cells decreases as *ρ*_1_ increases, while the density of spores is maximized for an intermediate value of *ρ*_1_. We will be specifically interested in the density of vegetative cells and spores that survive dispersal to colonize new demes. While we assume throughout the rest of this paper that only spores survive dispersal, our methods generalize to cases where there is a quantitative dispersal advantage of spores.

Figure 1 shows how the sporulation rate determines equilibrium spore density. Note that within a single patch the spore density is positive for a range of values of *ρ*, which always includes *ρ* near zero, so long as resource production is high enough relative to the vegetative cell death rate. Above a threshold value of *ρ*, the birth rate of vegetative cells is too low for the population to be sustained and both vegetative and sporulated cell density drops to 0.

**Figure 1.**
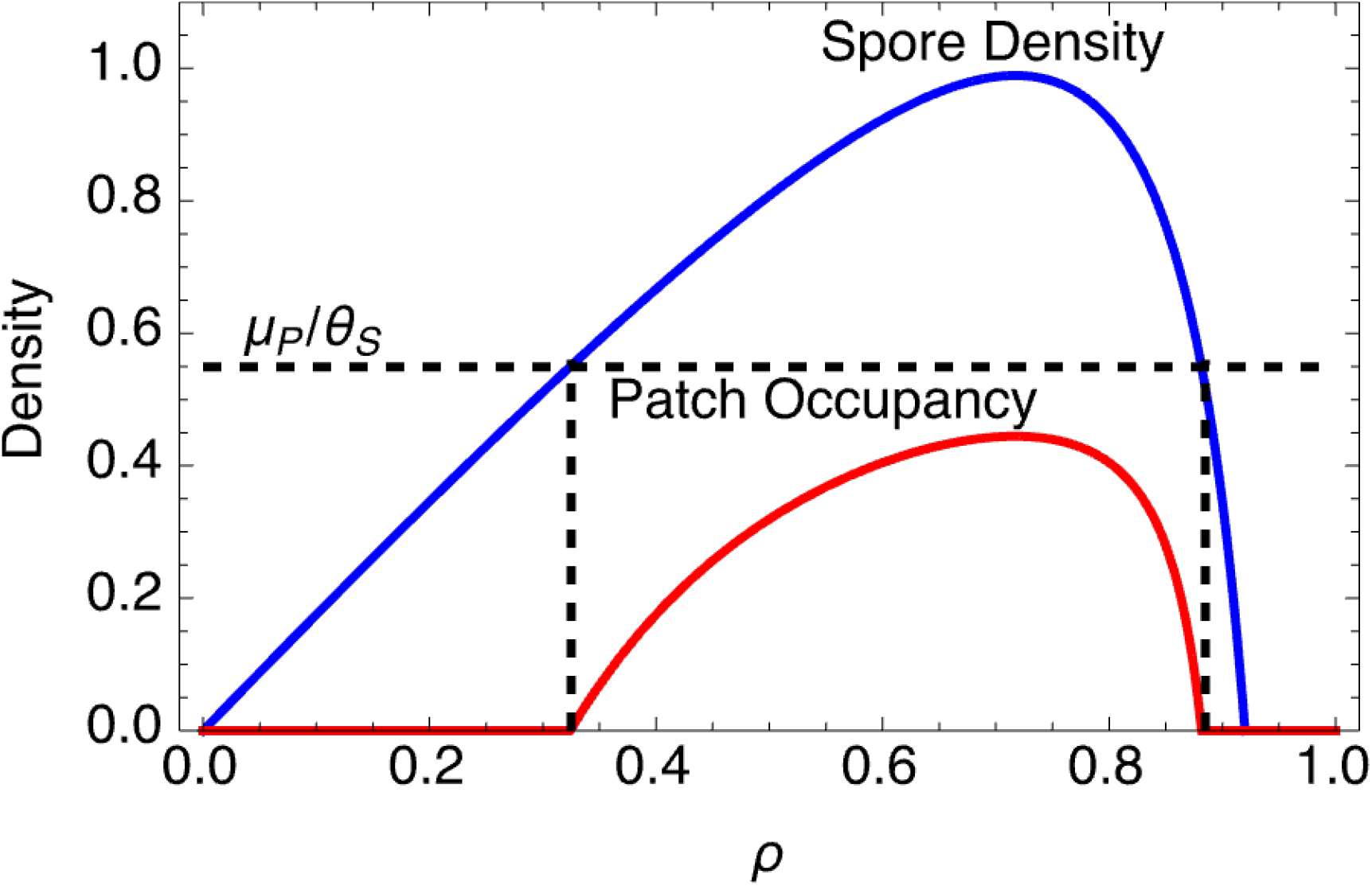
Spore density and metapopulation patch occupancy as a function of the rate of sporulation. Spore density, scaled as a fraction of the maximum, is shown in blue (the higher curve). Equilibrium patch occupancy is shown in red. Metapopulation persistence only occurs if the spore density is above the threshold *µ*_*P*_ */θ*_*S*_. Thus, if sporulation rate is too low, and spore density drops below the threshold, then the patch occupancy drops to zero. Conversly, if sporulation rate is too high, spore density drops below the threshold and patch occupancy drops to zero. Parameter values are *α* = 0.5, *µ*_*R*_ = 0.1, *µ*_*V*_ = 0.2, *µ*_*S*_ = 0.05, *r* = 1, *sc* = 1, *γ* = 0.5, *µ*_*P*_ = 0.5, *θ*_*S*_ = 0.35

### Metapopulation Dynamics

We assume that all patches achieve their within-patch equilibrium over a short time scale relative to metapopulation dynamics. This allows a much simpler analysis of the metapopulation level patch occupancy dynamics (we plan to consider non-equilibrium within-patch dynamics in future work).

We use the Tilman model of patch-occupancy (Tilman, 1982), based upon the Levins metapopulation model (Levins, 1969). There are *n* possible genotypes indexed in ascending order of their sporulation rates, the patch occupancy dynamics are

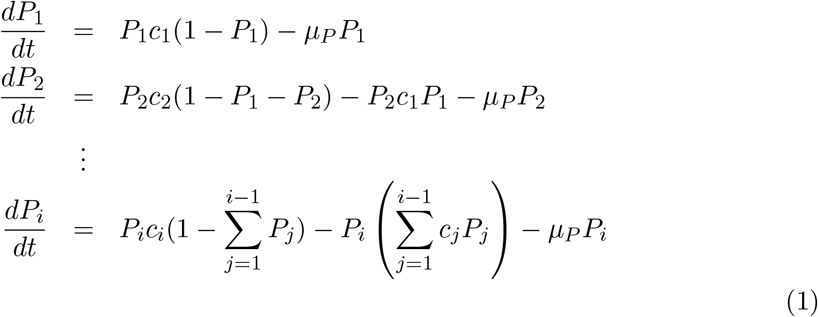

where *P*_*i*_ is the fraction of patches occupied by genotype 1, *c*_*i*_ is the colonization rate of genotype 1, and *µ*_*P*_ is the patch extinction rate. Note that the patch extinction rate is assumed to be independent of all within patch variables.

Based on our within-patch dynamic model, we define the colonization rate as

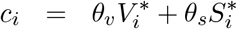

where the parameters *θ*_*i*_ represents the per-cell propensity of surviving dispersal and colonizing a new patch. For the present we assume that only spores are transmitted to new patches, such that *θ*_*v*_ = 0.

We can now calculate the equilibrium patch occupancy for a metapopulation containing a single genotype with the sporulation strategy *ρ* (figure 1). Note that metapopulation persistence only occurs for values of *ρ* in a specific range, and that patch occupancy will either remain constant or decrease when another genotypes is added.

Our first result is, given the parameters for resource dynamics, cell dynamics, and metapopulation dynamics, there is an intermediate range of *ρ* values that support population persistence. Also note that while within-patch dynamics show that low *ρ* is always favored, at the metapopulation level, patch occupancy is maximized at the same intermediate level of *ρ* that maximizes the equilibrium density of spores within a deme. If the value of *ρ* is too low the metapopulation will go extinct because colonization rates drop below the metapopulation persistence threshold, and if the value of *ρ* is too high, then within-deme extinction occurs because the birth rate is below the death rate.

Metapopulation persistence occurs if this condition is met:

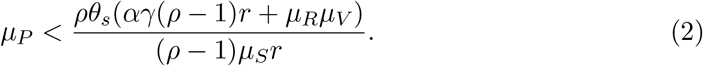

We define the minimum and maximum values of *ρ* that allow metapopulation persistence as *ρ*_min_ and *ρ*_max_ which can be found by rearranging the inequality (See supplemental Mathematica file).

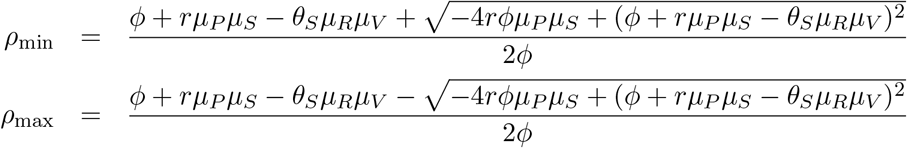

where *ϕ* = *rαγθ*_*S*_

### Invasion of Mutant Sporulation Strategies

We consider an invasion/replacement model of long-term evolution, as used in adaptive dynamics and weak mutation population genetics (Gillespie, 1991; Hammerstein, 1996; Dieckmann and Law, 1996; Champagnat et al., 2001; Proulx and Day, 2002; Lion, 2018). Given a resident genotype with sporulation rate *ρ*_1_ we can ask whether or not a mutant with a sporulation rate *ρ*_2_ can invade, and further whether that mutant will replace or coexist with the resident strategy. We consider a continuum of allele/strategy values in order to predict the sequence of population states following repeated rounds of mutation and selection.

This formulation follows from the Tilman (1982) model where the value of *ρ* places each strategy on the colonization/competition continuum. Lower values of *ρ* are better competitors, and the metapopulation frequency of a strategy is only affected by the presence and frequency of strategies with lower values of *ρ*. We can determine if a new sporulation strategy can invade given that the metapopulation is currently fixed for a single sporulation strategy 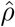 using the metapopulation invasion criteria. If the new sporulation strategy *ρ* is a lower sporulation rate, then it always has competitive advantage within a deme and will invade so long as *ρ*_min_ *< ρ < ρ*_max_. To determine if a strategy with 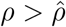 will invade we first solve for the equilibrium patch occupancy of the resident strategy, 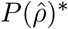 ^*∗*^, to find

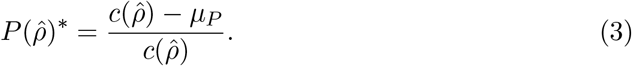

Invasion of a strategy with a higher sporulation rate can be determined by approximating around the invader being rare and inserting 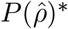 for the resident genotype to give

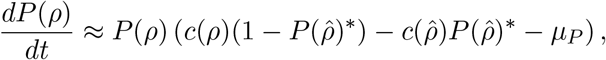

meaning that the mutant invades whenever

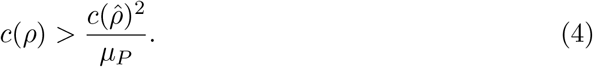

Note that *c*(*ρ*_min_) = *µ*_*P*_, and therefore if *c*(*ρ*) is an increasing function, at least for *ρ* near *ρ*_min_, then it is always the case that a small increase in *ρ* will lead to invasion and coexistence of the sporulation strategies.

We can use these calculations to define the possible scenarios for fate of a mutant sporulation strategy. For a mutant to invade, the mutant must be able to persist in the metapopulation even without competition from other strategies. If the mutant can persist on its own, it may also be able to invade the resident strategy, and then will either replace the resident strategy or coexist. A pairwise invisibility plot is shown in figure 2, where the colored regions show the outcome. The basic structure of this plot depends on the results that we have already laid out, and the size and position vary depending on the parameters (see supplemental Mathematica file for a dynamic version of the figure where parameters can be manipulated by the user).

**Figure 2.**
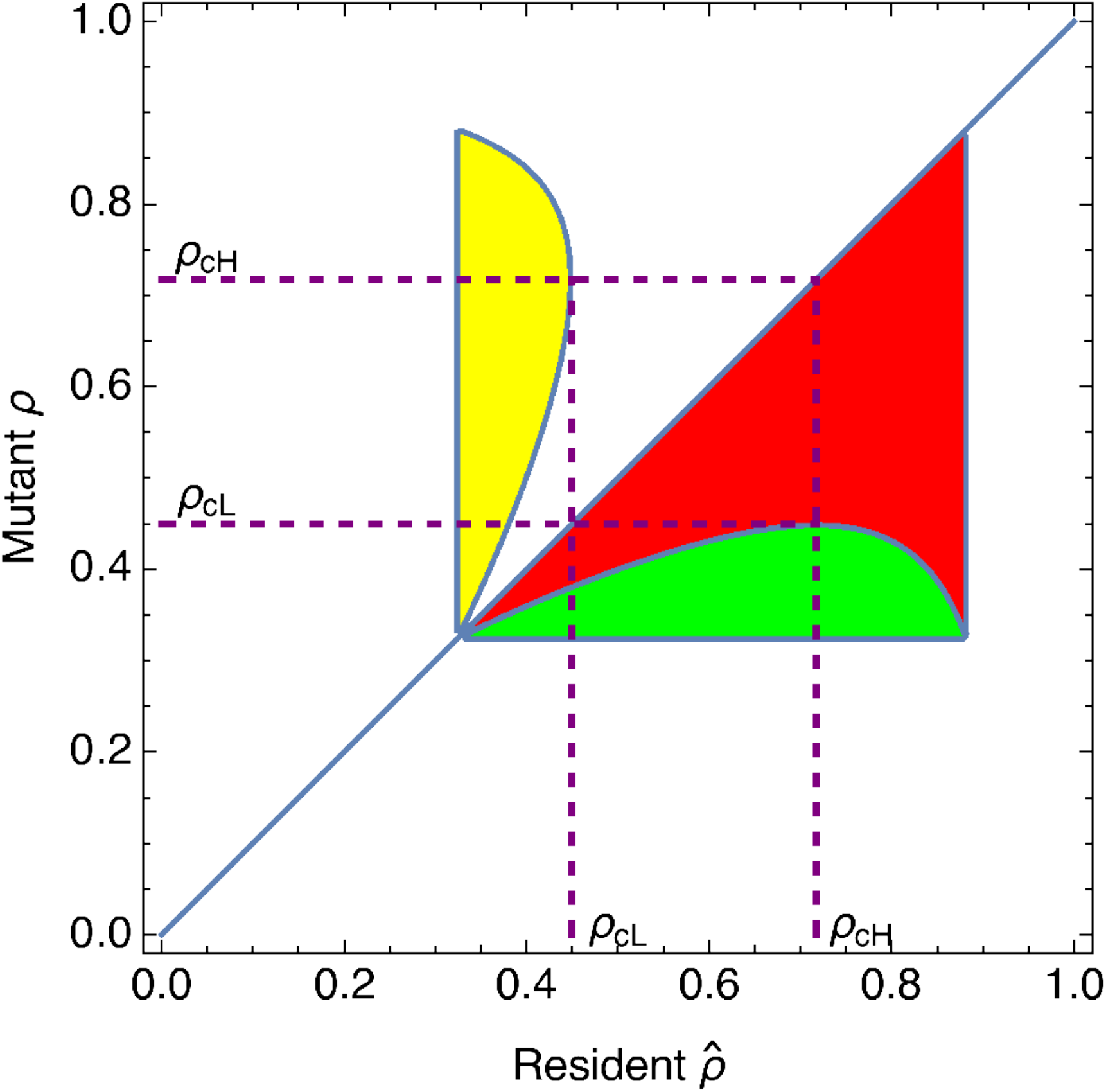
Pairwise invasibility plot. The red region shows areas where the mutant strategy can invade and would then replace the resident at the metapopulation level. The green and yellow areas both indicate that the mutant would invade but would then coexist with the resident at the metapopulation level. Parameter values are *α* = 0.5, *µ*_*R*_ = 0.1, *µ*_*V*_ = 0.2, *µ*_*S*_ = 0.05, *r* = 1, *sc* = 1, *γ* = 0.5, *µ*_*P*_ = 0.5, *θ*_*S*_ = 0.35

The pairwise invasibility plot shows first that only values of *ρ* satisfying *ρ*_min_ *< ρ < ρ*_max_ can invade, giving the outer square structure to the area. The red region indicates that the mutant strategy can invade and will replace the resident (i.e. the two strategies do not coexist). Both the green and yellow regions shows that the mutant can invade but will coexist with the resident. The distinction between the green and yellow region is that in the yellow region the mutant has a higher sporulation rate, and in the green region it has a lower sporulation rate.

The pairwise invasibility plot can be used to describe the long-term evolutionary dynamics. If the resident strategy is above *ρ*_min_ then mutants that have a small decrease in *ρ* can invade and replace the resident. Under the typical weak mutation model (Gillespie, 1991; Lion, 2018) we would expect repeated rounds of mutation and fixation to result in a decrease in *ρ* until it gets near to *ρ*_min_. In this area, where the three colors are all near each other, mutations that increase *ρ* enough can then both invade and coexist. Regardless of the specific parameters, we always expect coexistence of at least two sporulation strategies.

Examining the shape of the yellow region, we see that for values of *ρ* near *ρ*_min_, small increases in *ρ* can invade, and as the resident value of *ρ* increases, larger increases in *ρ* are required to allow invasion. The range of mutant *ρ* values that can invade shrinks as *ρ* continues to increase, and finally is narrowed down to a single point. We name the resident value *ρ*_cL_ for the largest value that can be invaded by a larger value of *ρ*. We call the value of *ρ*_cH_ the value of *ρ* that can invade *ρ*_cL_. Figure 2 shows how these values can be found graphically. We see that for a resident population at *ρ*_cH_ the range of values that can invade and coexist are the values between *ρ*_min_ and *ρ*_cL_.

Although the pairwise invasibility plot does not allow for us to graphically determine how more than two strategies can coexist, we believe it implies that coexisting strategies will be restricted to *ρ < ρ*_cH_. This would occur because initially lower values of of *ρ* would invade until the population entered the region between *ρ*_cL_ and *ρ*_cH_. At this point, mutations that produced local increases in *ρ* could invade and coexist. Once the largest value of *ρ* is near to *ρ*_cH_ further mutations that increase *ρ* will not invade.

### Coexistence of a continuously distributed set of sporulation strategies

Using our pairwise invasibility analysis, we were able to show that a range of strategies can coexist. However, the PIP approach does not allow for us to calculate invasibility of additional mutants when two genotypes are already coexisting. To address this issue, we developed another technique that models the population density of genotypes as a continuous frequency distribution. From this we derive an integro differential equation for the change in frequency distribution and find an equilibrium distribution that represents an ESS containing a continuum of competing strategies.

We use the variable *ρ* to represent the sporulation strategy. The density is defined as *P* (*ρ*)_*t*_ at time *t*. We start by defining a discrete time updating process where the density at time *t* is given by

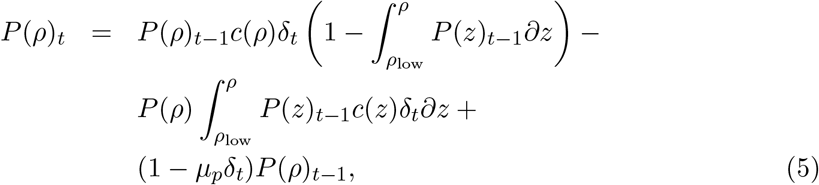

which captures the increase of density due to occupying patches that are currently occupied by weaker competitors, the loss of density to better competitors, and the continued occupancy of patches that have not gone extinct due deme extinction. Here we set the minimum genotype present to *ρ*_low_. Standard rearrangement and taking the limit as *δ*_*t*_ *→* 0 gives

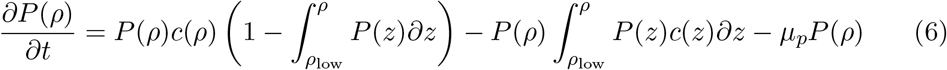

The steady state can be solved for by setting

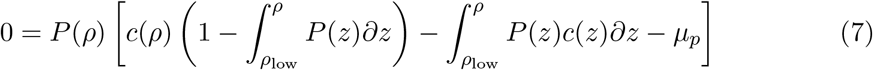

which holds if either *P* (*ρ*) = 0 for all *ρ* or the term inside the square brackets is 0.

Focusing on the second (i.e. non-trivial) equilibrium from equation 7, we can differentiate with respect to *ρ* to get an ODE that describes the steady state.

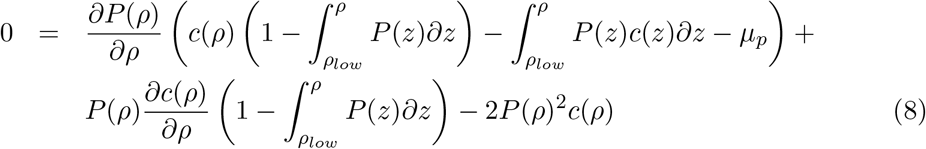

Substituting equation 7 into equation 8 we have

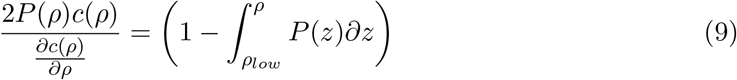

Taking the derivative with respect to *ρ* now yields an ODE

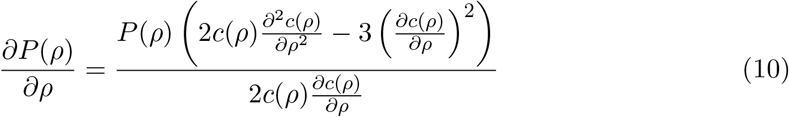

Our function *c*(*ρ*) is given from our model for the within-patch dynamics. This differs from other models of species packing, where the trade-off between competition and colonization is more abstract (Kinzig et al., 1999). The differential equation can be solved analytically and leaves us with an additional constant of integration that must be set based on ecological assumptions (see supplemental Mathematica file for the solution of this ODE). The value of the constant of integration is determined by our assumption that there is a lower bound on the sporulation strategies included in the solution. We solve for the density of this lowest sporulation rate strain and use that to determine the constant of integration.

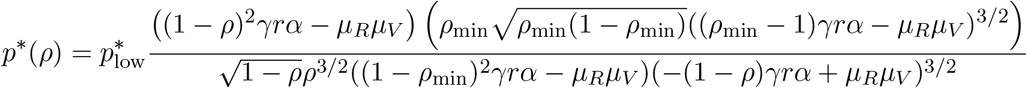

Where *ρ*_min_ is the minimum value of *ρ* included in the set of coexisting genotypes, and 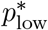 represents the equilibrium patch occupancy frequency of the genotype with the lowest sporulation rate. Because 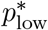 appears as an independent factor multiplying the other terms, changing 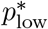 does not alter the value of *ρ* where the patch occupancy frequency reaches 0, i.e. the highest sporulation rate that can be included in the coalition of strategies.

The first term inside the parentheses in the numerator is the only term in the numerator that involves *ρ*, and therefore the sign of this term determines the sign of the numerator. It represents a balance between the resource production and consumption terms and the resource and cell clearance rates.

Solving for the when the sporulation rate specific term is greater than zero gives the critical value of *ρ* above which the patch occupancy rate is zero. This is

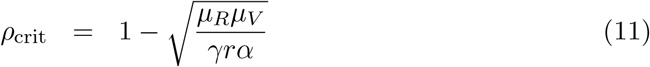

We can substitute our result from this section into an expression for the boundary of the green region in the PIP plot. Taking the derivative of that curve and substituting in the value of *ρ*_crit_ yields 0, showing that this point coincides with the extremum of the boundary of green region. This means that values of the area of coexistence can be deduced from the PIP and includes all values of *ρ* between *ρ*_cH_ and the thickest part of the green region.

Figure 3 shows how the density of strategies varies with *ρ* and depends on the patch extinction rate (*µ*_*P*_). The upper threshold value for coexistence is independent of *µ*_*P*_, but the minimum level of sporulation to maintain the metapopulation does depend on *µ*_*P*_. The relative density is largely independent of the patch extinction rate, so that the curves are very similar. Higher patch extinction rates lead to a smaller range of coexisting *ρ* values, and also have lower total patch occupancy levels. We calculated the total patch occupancy by integrating the density of strategies over the range of *ρ* that allow coexistence/metapopulation persistence.

**Figure 3.**
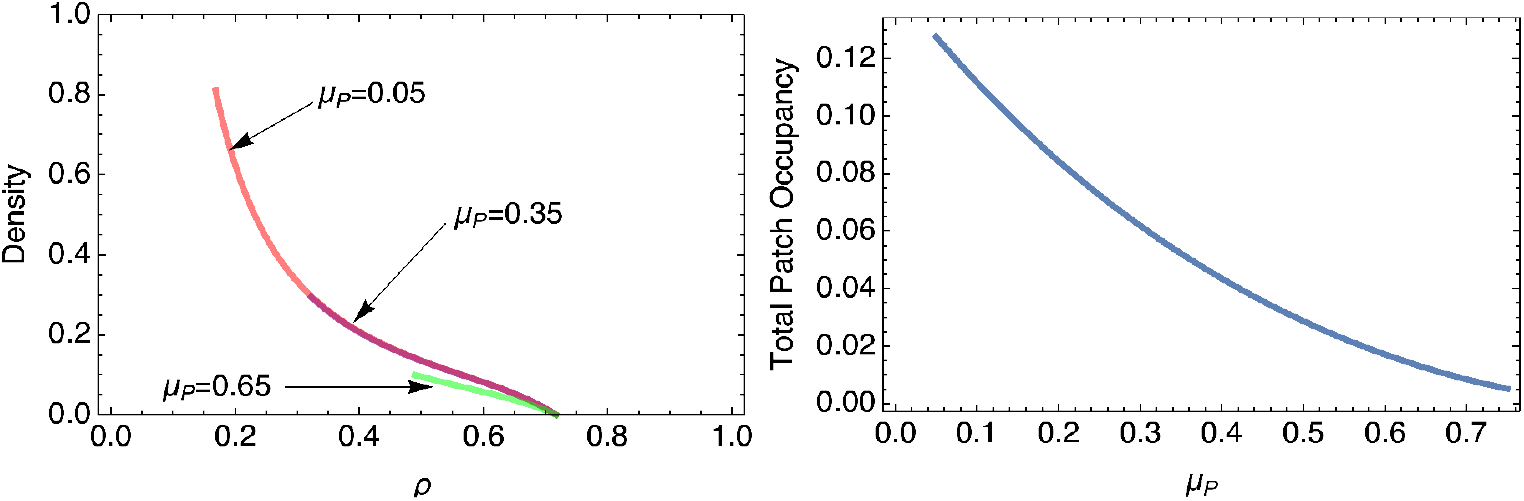
Plots of the density of coexisting sporulation strategies. Panel A shows the density of different *ρ* strategies. The value of *µ*_*P*_ is varied and indicated for each curve. Panel B shows the total patch occupancy as a function of the patch extinction rate. Parameter values are *α* = 0.5, *µ*_*R*_ = 0.1, *µ*_*V*_ = 0.2, *µ*_*S*_ = 0.05, *r* = 1, *sc* = 1, *γ* = 0.5, and *θ*_*S*_ = 0.35

## Discussion

Spore formation in yeasts has often been described as an adaptation to survive environmental conditions not favorable to cell growth, and has more recently been viewed as an adaptation to increase survival while being vectored by insects, i.e. phoretic dispersal (Coluccio et al., 2008; Reuter et al., 2007), and has also been discussed as a form of kin selection (Ratcliff et al., 2013). Recently, Twyman and Gardner (2023) have presented a very general analysis of the evolution of dormancy where the strength of selection on increased dormancy can be calculated based on knowledge of the degrees of relatedness among various groups in a metapopulation.

We developed our model to ask how the dynamics of competition within patches and colonization between patches might lead to the evolution of sporulation. We developed our framework by considering a simple chemostat type dynamic of resource renewal within demes and dispersal between demes via quiescent spores. We took two approaches to understanding the long-term evolution of sporulation, one based on pairwise-invasibility and the other by modeling competition among a continuous distribution of trait values. Similar to other work on the competition-colonization model, we found that genotypes of higher within-deme competitiveness excluded a range of genotypes with higher dispersal abilities (Kinzig et al., 1999). However, the specific dynamics that we modelled allow us to predict a restricted range of possibly coexisting strategies. We predict that recurrent mutations of small effect initially result in a decline in sporulation rate. Subsequently, as the resident genotype approaches the minimum sporulation rate necessary for a sustainable metapopulation, mutations with more substantial effects promoting increased sporulation rate can coexist.

When we extended our analysis to include a continuum of competing strategies we found that a continuous set of sporulation strategies can coexist. The strategies that show a trade-off between within deme competition and dispersal coexist, but we also found that above a critical rate of sporulation, equilibrium spore density begins to decrease as sporulation rate increases. In other words, above this critical point *ρ*_*cH*_, strategies with higher sporulation are both weaker dispersers and weaker competitors than some lower *ρ* strategies that are part of the coexisting set of genotypes.

We assumed that yeast have a fixed strategy in terms of when they produce spores. Yeast sporulation tends to be triggered by chemical conditions that correlate with depletion of sugars and crowding of yeast (Neiman, 2005). Natural yeast isolates do vary in their propensity to sporulate (Liti et al., 2009), and vary in their ability to evolve increased sporulation rates (Thomasson et al., 2021). Less is known about how sporulation proceeds under natural conditions and to what extent sporulation represents a facultative program. In the context of our model, we might expect a strategy that only grows vegetatively until resources are depleted to their equilibrium level. However, this would not change the results of within-deme competition so long as both strains survive until resource levels approach their equilibrium. After that point, both strains would produce spores at their genotype-specific rates, the strain with lower sporulation rate would come to dominate. Likewise, strains who colonize a patch alone would reach their population dynamic equilibrium more quickly in the absence of sporulation, and then begin sporulating and reach their spore-density equilibrium.

When we considered pairwise invasion, we found that strategies with lower sporulation rates can invade those with higher rates, and that this can continue to proceed until the strategy with the minimum sporulation rate for a viable metapopulation has invaded. Paradoxically, this ”minimum viable” sporulation strategy has an equilibrium metapopulation density of 0. Clearly, a strategy with a metapopulation density of 0 does not really exist. One way that we could extend the model to reconcile this is by incorporating the probability of fixation in our calculation for whether a mutant strategy will invade. We expect that including this would allow strategies near the minimum viable sporulation rate to be excluded. For our continuous strategy distribution model, we did assume a lower bound of on sporulation rate, which is effectively assuming that strategies below that sporulation rate are unable to persist.

One criticism of Tilman (1994)’s competition-colonization trade-off model is that it assumes competition to deterministic and to have a binary outcome. In particular, the magnitude of a strategies competitive advantage does not alter the degree of the outcome (Adler and Mosquera, 2000). This follows from the assumption that within-deme dynamics play out over a long enough time with the competitively superior type reaching an equilibrium density that is not at all affected by the competitor. Several relevant ecological processes could alter this assumption and deserve further consideration.

For yeast, two relevant ecological scenarios that could alter this assumption include situations where the dispersing insects visit patches before they reach equilibrium and situations where the within deme dynamics are not chemostat-like. There is evidence that flies are attracted to yeast products, and so we might expect only mature patches to be visited. In general, however, flies could visit patches at any time, not only once they have neared equilibrium. It may be possible to approximate this effect by including the average lifespan of patches and approximating the rate of approach to equilibrium. We suspect that this would have little effect on our results unless that rate of approach to the equilibrium decreases steeply as we get closer to an evolutionary attracting point. The other relevant ecological scenario is one where competition within demes follows from a process where equilibrium densities are not independent of initial conditions. For example, a resource pulse model, similar to serial transfer, would have quite different dynamics. Under such a scenario, competitors would extract resources that would not be renewed, and would therefore affect the total amount of cells and spores that other strains would be able to produce.

In this analysis, we find that multiple sporulation strategies can coexist, and they do so because a trade-off between spore production and within-deme competitive ability naturally arises. Another scenario in which multiple dormancy behaviors coexist is when seeds can decide whether to germinate or to remain dormant. When the environmental conditions that germinating seeds face vary from generation to generation, then a randomizing strategy can be favored because it increases the geometric mean of within-generation fitness (Cohen, 1966; Bulmer, 1984; Ellner, 1987; Philippi and Seger, 1989). In these cases, polymorphism in the propensity to germinate cannot be maintained as genetically based differences, but rather as phenotypically variance that maximized geometric mean fitness. In particular, this would not lead to coexistence of distinct species that vary in their germination strategy.

The yeasts *S. cerevisiae* and *S. paradoxus* are closely related, coexist in some natural habitats, and vary in their sporulation and germination programs (Sniegowski et al., 2002; Murphy and Zeyl, 2012). The observed disparities in temperature sensitivity do not provide an explanation for their coexistence. Our theoretical framework shows that the divergence in sporulation strategies, even in the absence of discernible distinctions in other physiological characteristics, may serve as a pivotal factor contributing to the coexistence of these yeast species. Further empirical validation of this hypothesis would advance our understanding of microbial coexistence and contribute to the broader comprehension of ecological interactions within microbial communities.

## Supporting information

Supplemental Mathematica Code

## A Within Patch Stable Equilibrium

Here we prove that the yeast strain with lowest sporulation rate becomes fixed when we model dynamics within a single patch. This can be thought as a case of the competitive exclusion principle. The system of ODEs is:

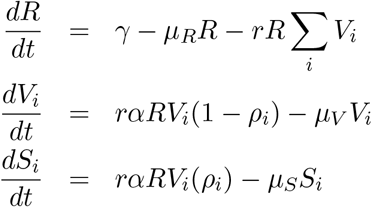

We assume that all strains start with a non-zero vegetative cell population.

### Result 1

Say there are *n* total yeast strains each with a unique sporulation rate *ρ*_*i*_. We index the strains so that *ρ*_1_ *< ρ*_2_ *<* … *ρ*_*n*_. Then there are two types of equlibria. The first is where all strains are extinct and only resources are present.

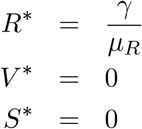

This equilibrium is stable if no strain is fit enough to survive, which happens if and only if 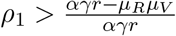 Otherwise if 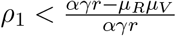 then for each strain *i* we have the equilibria

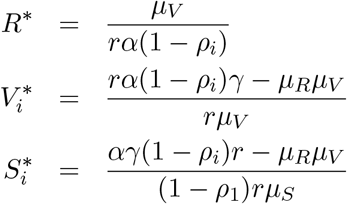

where all the other strains are at zero. Of these equilibria the only stable one is that where the best competitor, strain 1, is fixed.

*Proof*. To begin observe that the equations for the spores do not influence the resources or vegetative cells. Their behavior is entirely determined by the dynamics of vegetative cells and the resource. This means for most of the analysis we can ignore the spores and focus on the vegetative/resource subsystem.

### Part 1: These are the only equilibrium

If the equilibrium contains no vegetative cells then all spores eventually decay to zero and a quick calculation shows that the resource equilibrium is *R*^*∗*^ = *γ/µ*_*R*_ Otherwise say that we are at an equilibrium where the *i*th strain is non-zero. This strain implies a unique value for *R*^*∗*^.

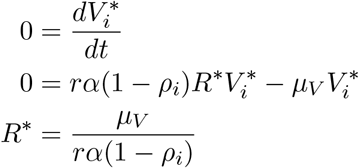

Each *ρ*_*i*_ is unique, meaning two strains cannot coexist as they would each imply two different values for *R*^*∗*^. Thus the *i*th strain is the only strain present, which simplifies the equation 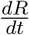. Substituting *R*^*∗*^ into this equation gives the equilibrium for strain *i*.

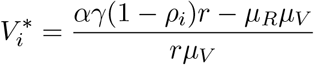

This in turn is all we need to know to calculate the equilibrum for the spores.

Observe that the sporulated cell densities do not influence the resource-vegetative dynamics. Rather, spores follow the dynamics of the resources and vegetative cell subsystem. The stability of the equilibria do not change when remove the spore equations from the system. We do this for the rest of the analysis.

### Part 2: The equilibrium with strain 1 is stable as long as 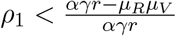

We show this by standard Jacobian stability analysis. Let *ω*_*i*_ = *rα*(1 *−ρ*_*i*_). The Jacobian matrix for this system is then

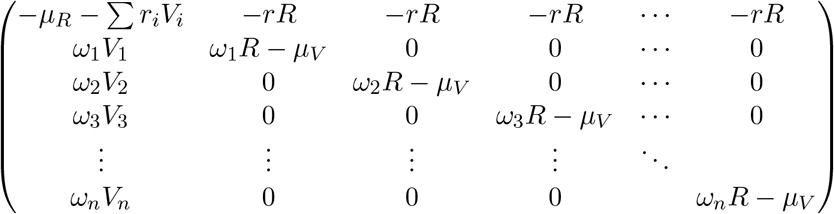

We will look at this matrix around the equilibrium where strain 1 is fixed. Written in terms of *ω*_1_ the vegetative equilibrium is 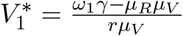 and the resource equilibrium 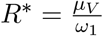 Substituting these in give

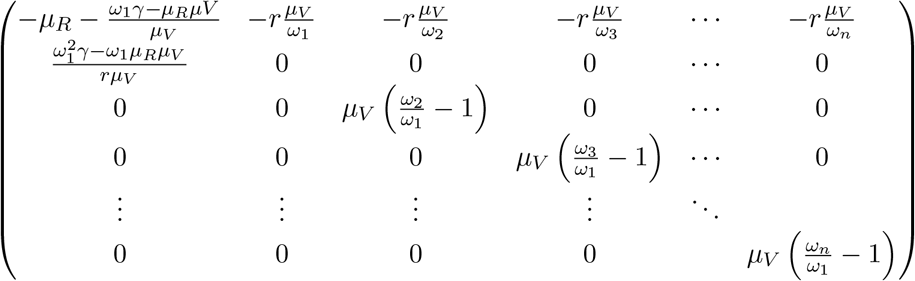

The first row of this matrix must be negative, as all the *ω*_*i*_ are positive. Furthermore the remaining terms on the diagonal must be negative as *ω*_1_ is strictly larger than the other *ω*_*i*_ (as *ρ*_1_ is the smallest). The signs of this matrix look like.

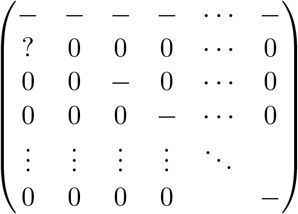

The question mark entry is the only one of unknown sign, that of 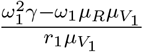

To find the eigenvalues of this matrix we can view it as an adjacency matrix. Then it corresponds to a network with a simple structure. The resource vertex is strongly connected with the strain 1 vertex. The remaining strains are strongly connected only with themselves via a self-loop, giving a total of *n −* 1 trivial connected components.

A well known result from spectral graph theory says that the strongly connected components of a network give the eigenvalues. The single node components *V*_2_ through *V*_*n*_ each give negative eigenvalues as their self loops lie on the diagonal of the Jacobian matrix. The two node component strongly connecting *V*_1_ and *R* contributes the same eigenvalue as the matrix

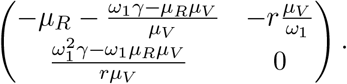

We calculate that this matrix has only one eigenvalue,

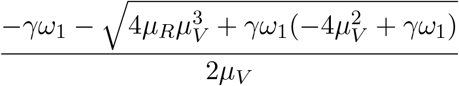

which a little algebra shows is negative if and only if 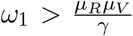 Rewriting this in terms of *ρ*_1_ we find the condition for all eigenvalues of the Jacobian to be negative is

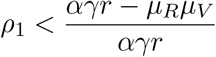

This is the same condition as the resource only equilibrium being unstable. Said another way, the sporulation rate is low enough that the vegetative yeast is able to survive. In this case the strain with lowest sporulation rate is always stable as it is the best competitor. If we start a patch with a set of strains the one with the lowest sporulation rate will take over.

### Part 3: All other equilibria with a strain present are unstable

Take the Jacobian matrix and evaluate it at the equilibrium where the *i*th strain is present. Without loss of generality say we evaluate at strain 2. Then the Jacobian is

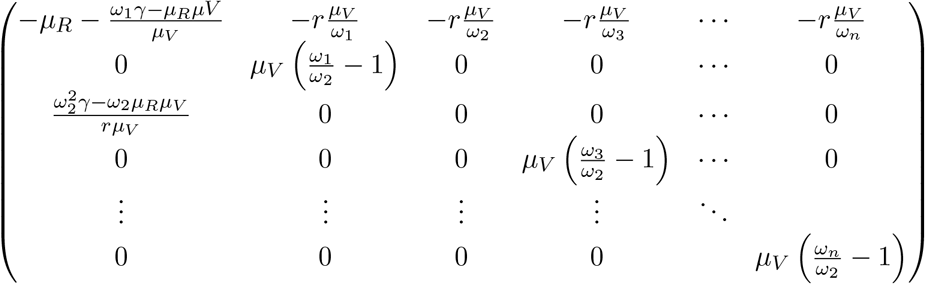

Like before, if we consider this as an adjacency matrix then it is made up of many strongly connected components. Most of these are trivial, consisting of only one node. In particular the node corresponding to *V*_1_ is one of these trivial components. It has only an edge pointing out of it to the resource node and a self loop with value 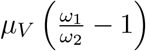 This self loop value is the eigenvalue and must be positive. Thus this equilibrium is unstable.

### Part 3: The resource only equilibrium is stable equilibrium if 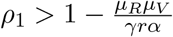

Next we will look at the equilibrium where only resources are available. Setting *R* to the equilibrium value *R*^*∗*^ = *γ/µ*_*R*_ and all of the *V*_*i*_ to zero means the Jacobian becomes

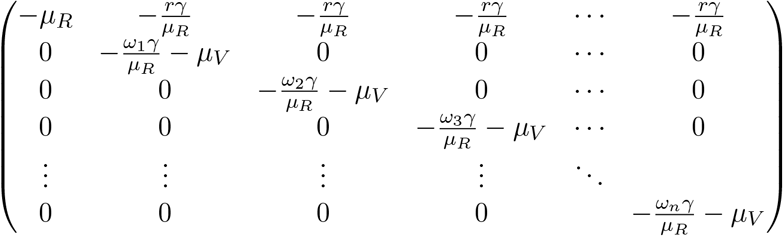

The eigenvalues lie on the diagonal as this matrix is upper triangular. The first term of the diagonal is obviously negative, but what about the others? Under what conditions are they all negative, implying stability? Solving for *−ω*_*i*_*γ/µ*_*R*_ *−µ*_*V*_ *<* 0 and substituting in the value for *ω*_*i*_ = *rα*(1 *− ρ*_*i*_) we find the condition is

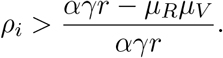

If the *i*th strain meets this condition it means it makes so many spores it cannot survive in the environment, regardless of competition. Not enough of its resources go to reproduction. This is automatically true for all strains when *i* = 1 in the above equation. When the inequality sign is flipped then resources only is not a stable equilibrium and strain 1 fixes.

When *ω*_*i*_ is seen as fitness this result can be considered a case of the competitive exclusion principle. A similar arguement can extend these results to the case where each strain has a unique growth rate and death rate (with the caveat that the ratio of fitness and death must be unique, that is *µ*_*Vi*_ *≠*_*i*_ ≠ *µ*_*Vj*_ *≠*_*j*_).

This proof does not consider limit cycles and chaotic dynamics. This is in part because there is no known general way to find these for high dimensional systems such as this one. But we also do not see any trace of these in numerical calculations, and such cycles and chaos tend not to exist for chemostat-style models such as this.

