## Supplemental Mathematica Code for "Selection on sporulation strategies in a metapopulation can lead to coexistence"

### Figures for theory paper on the evolution of sporulation in yeast

In[287]:=

```
SetDirectory[
 "/Users/proulx/Dropbox/Apps/Overleaf/Evolution of Sporulation/Figures"]
```

Out[287]=

```
/Users/proulx/Dropbox/Apps/Overleaf/Evolution of Sporulation/Figures
```

A strain producing $\rho$ fraction of spores per division will have an equilibrium density of spores that is maximal for an intermediate $\rho$.

In[ ]:=

$$\text{Plot}\left[\frac{\rho\,(-r\,\alpha\,\gamma+\rho\,r\,\alpha\,\gamma+\mu R\,\mu V)}{(-1+\rho)\,r\,\mu S}\;/.\;\{\alpha\to .5,\ \mu R\to 0.1,\ \mu V\to .2,\ \mu S\to .05,\ r\to 1,\ \theta S\to 1,\ \gamma\to 0.5,\ \mu P\to 0.5\},\ \{\rho,\ 0,\ 1\}\right]$$

Out[ ]=

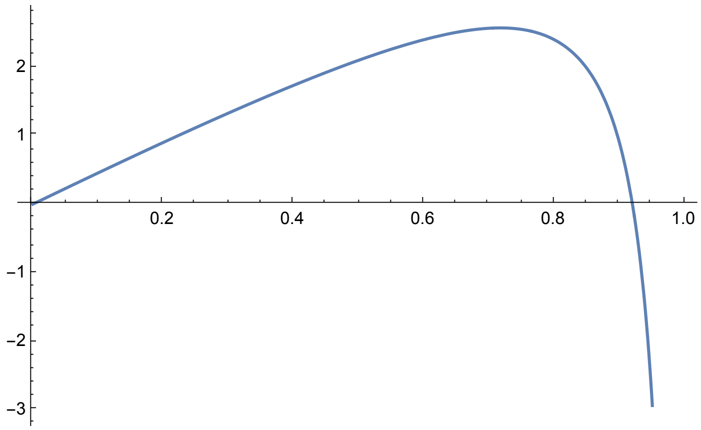

In[ ]:=
$$\text{SporeDensityPlot = Plot}\left[\frac{\rho\,(-r\,\alpha\,\gamma+\rho\,r\,\alpha\,\gamma+\mu R\,\mu V)}{(-1+\rho)\,r\,\mu S}\;/.\right.$$

```
  {α → .5, μR → 0.1, μV → .2, μS → .05, r → 1, θS → 1, γ → 0.5, μP → 0.5},
  {ρ, 0, 1}, PlotRange → {0, 3}, PlotStyle → {Black, Thick},
  Frame → True,
  FrameLabel → {"ρ", "Spore Density"}, BaseStyle →
   {FontSize → 12, FontWeight → Plain, FontFamily → "Helvetica"}, ImageSize → 300]
```

Out[ ]=

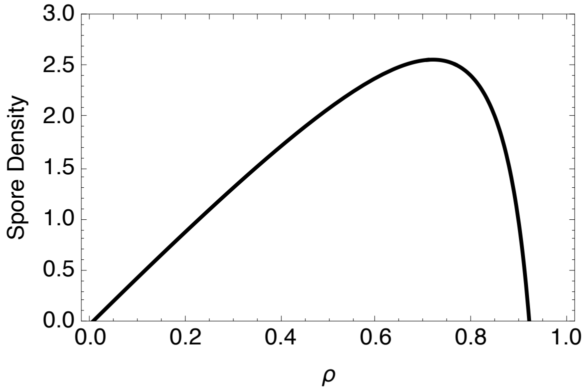


In[ ]:=
```
Export["SpoDensityPlot.pdf", Show[SporeDensityPlot,
  Prolog → {Opacity[0], Texture[{{{0, 0, 0, 0}}}], VertexTextureCoordinates →
     {{0, 0}, {1, 0}, {1, 1}}, Polygon[{{0, 0}, {.1, 0}, {.1, .1}}]
   }]]
```

Out[ ]=

SpoDensityPlot.pdf

The equilibrium patch occupancy for a single strain with sporulation rate $\rho$ is

In[ ]:=
$$\text{Simplify}\left[\text{Solve}\left[\theta S * \frac{\rho\,(-r\,\alpha\,\gamma+\rho\,r\,\alpha\,\gamma+\mu R\,\mu V)}{(-1+\rho)\,r\,\mu S}\,P\,(1-P)-\mu P\,P == 0,\ P\right]\right]$$

Out[ ]=
$$\left\{\{P\to 0\},\ \left\{P\to \frac{\theta S\,\mu R\,\mu V\,\rho + r\,(-1+\rho)\,(-\mu P\,\mu S+\alpha\,\gamma\,\theta S\,\rho)}{\theta S\,(\mu R\,\mu V + r\,\alpha\,\gamma\,(-1+\rho))\,\rho}\right\}\right\}$$

So the condition for metapopulation persistence is given by

In[ ]:=
$$\text{Simplify}\left[\text{Solve}\left[\frac{\rho^2\,r\,\theta S\,\alpha\,\gamma + r\,\mu P\,\mu S - \rho\,(r\,\theta S\,\alpha\,\gamma + r\,\mu P\,\mu S - \theta S\,\mu R\,\mu V)}{\rho\,\theta S\,((-1+\rho)\,r\,\alpha\,\gamma+\mu R\,\mu V)} == 0,\ \mu P\right]\right]$$

Out[ ]=
$$\left\{\left\{\mu P\to \frac{\theta S\,(\mu R\,\mu V + r\,\alpha\,\gamma\,(-1+\rho))\,\rho}{r\,\mu S\,(-1+\rho)}\right\}\right\}$$

The minimum and maximum values for $\rho$ are given by:

In[ ]:=

$$\text{Simplify}\left[\text{Solve}\left[\frac{\rho^2\ r\ \theta S\ \alpha\ \gamma + r\ \mu P\ \mu S - \rho\ (r\ \theta S\ \alpha\ \gamma + r\ \mu P\ \mu S - \theta S\ \mu R\ \mu V)}{\rho\ \theta S\ ((-1+\rho)\ r\ \alpha\ \gamma + \mu R\ \mu V)} == 0,\ \rho\right]\right]$$

Out[ ]=

$$\left\{\left\{\rho \to \frac{r\ \alpha\ \gamma\ \theta S + r\ \mu P\ \mu S - \theta S\ \mu R\ \mu V - \sqrt{-4\ r^2\ \alpha\ \gamma\ \theta S\ \mu P\ \mu S + (r\ \alpha\ \gamma\ \theta S + r\ \mu P\ \mu S - \theta S\ \mu R\ \mu V)^2}}{2\ r\ \alpha\ \gamma\ \theta S}\right\},\right.$$
$$\left.\left\{\rho \to \frac{r\ \alpha\ \gamma\ \theta S + r\ \mu P\ \mu S - \theta S\ \mu R\ \mu V + \sqrt{-4\ r^2\ \alpha\ \gamma\ \theta S\ \mu P\ \mu S + (r\ \alpha\ \gamma\ \theta S + r\ \mu P\ \mu S - \theta S\ \mu R\ \mu V)^2}}{2\ r\ \alpha\ \gamma\ \theta S}\right\}\right\}$$

In[ ]:=

$$\text{PatchOccPlot} = \text{Plot}\left[\left(\frac{\rho^2\ r\ \theta S\ \alpha\ \gamma + r\ \mu P\ \mu S - \rho\ (r\ \theta S\ \alpha\ \gamma + r\ \mu P\ \mu S - \theta S\ \mu R\ \mu V)}{\rho\ \theta S\ ((-1+\rho)\ r\ \alpha\ \gamma + \mu R\ \mu V)}\right) * \text{If}\left[\left(\frac{\rho\ \theta S\ ((-1+\rho)\ r\ \alpha\ \gamma + \mu R\ \mu V)}{(-1+\rho)\ r\ \mu S} > \mu P\right), 1, 0\right]\ /.\right.$$

```
 {α → .5, μR → 0.1, μV → .2, μS → .05, θS → 0.35, γ → .5, μP → 0.5, r → 1},
 {ρ, 0, 1}, PlotRange → {0, 0.5}, PlotStyle → {Black, Thick},
 Frame → True,
 FrameLabel → {"ρ", "Patch Occupancy"}, BaseStyle →
  {FontSize → 12, FontWeight → Plain, FontFamily → "Helvetica"}, ImageSize → 300]
```

Out[ ]=

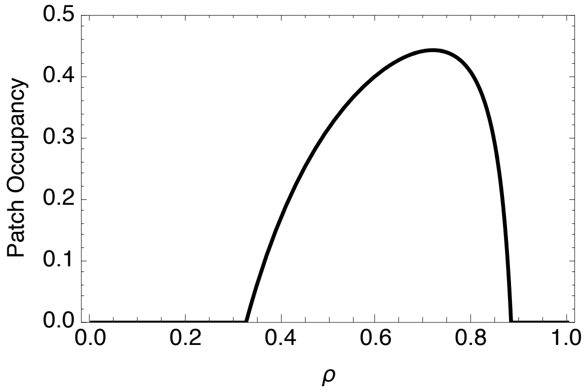


In[ ]:=

```
Export["PatchOccPlot.pdf", Show[PatchOccPlot,
  Prolog → {Opacity[0], Texture[{{{0, 0, 0, 0}}}], VertexTextureCoordinates →
     {{0, 0}, {1, 0}, {1, 1}}, Polygon[{{0, 0}, {.1, 0}, {.1, .1}}]
   }]]
```

Out[ ]=

```
PatchOccPlot.pdf
```

A figure that combines the sporulation density and patch occupancy density. We rescale the height of the spore density to be relative to the maximum.

In[289]:=

```
SporeDensityPlot = Plot[Evaluate[{ρ (-r α γ + ρ r α γ + μR μV) / ((-1 + ρ) r μS), μP / θS} / 2.6 /.
    {α → .5, μR → 0.1, μV → .2, μS → .05, r → 1, θS → 0.35, γ → 0.5, μP → 0.5, r → 1}],
  {ρ, 0, 1}, PlotRange → {0, 1.1}, PlotStyle → {{Blue, Thick}, {Black, Dashed}},
  Frame → True,
  FrameLabel → {"ρ", "Density"}, BaseStyle →
   {FontSize → 12, FontWeight → Plain, FontFamily → "Helvetica"}, ImageSize → 300];
```

```
PatchOccPlot = Plot[((ρ^2 r θS α γ + r μP μS - ρ (r θS α γ + r μP μS - θS μR μV)) / (ρ θS ((-1 + ρ) r α γ + μR μV))) *
    If[((ρ θS ((-1 + ρ) r α γ + μR μV)) / ((-1 + ρ) r μS) > μP), 1, 0] /.
   {α → .5, μR → 0.1, μV → .2, μS → .05, θS → 0.35, γ → .5, μP → 0.5, r → 1},
  {ρ, 0, 1}, PlotRange → {0, 0.5}, PlotStyle → {Red, Thick},
  Frame → True,
  FrameLabel → {"ρ", "Patch Occupancy"}, BaseStyle →
   {FontSize → 12, FontWeight → Plain, FontFamily → "Helvetica"}, ImageSize → 300];
```

```
ComboPlot = Show[SporeDensityPlot, PatchOccPlot, Epilog → {Directive[lineStyle],
    Line[{{.325, 0}, {.325, .56}}], Line[{{.885, 0}, {.885, .56}}],
    Text["μP/θS", {0.1, .6}],
    Text["Spore Density", {0.7, 1.03}],
    Text["Patch Occupancy", {0.55, .49}] }]
```

Out[291]=

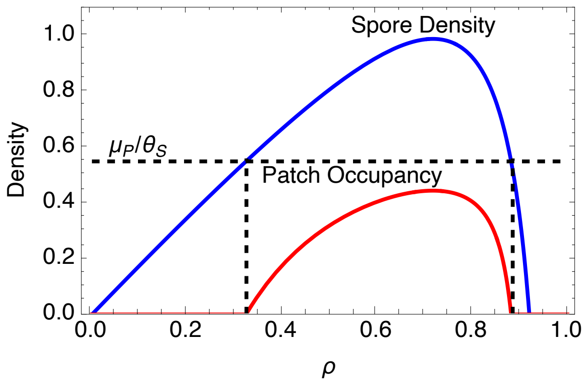

In[292]:=

```
Export["ComboPlot.pdf", Show[ComboPlot,
  Prolog → {Opacity[0], Texture[{{{0, 0, 0, 0}}}], VertexTextureCoordinates →
     {{0, 0}, {1, 0}, {1, 1}}, Polygon[{{0, 0}, {.1, 0}, {.1, .1}}]
   }]]
```

Out[292]=

```
ComboPlot.pdf
```

Create the PIP plots

In[ ]:=
```
spoInvPlot = Show[
  RegionPlot[(θS (k2 (-r α γ + k2 r α γ + μR μV))/((-1 + k2) r μS) > (θS (k1 (-r α γ + k1 r α γ + μR μV))/((-1 + k1) r μS))^2 / μP
     (*k2 has a sporulation advantage that is large enough to allow
      coexistence*) && k2 > k1 (*Just need to make sure that k1 is
      actually the better competitor*) && k1 < (r α γ - μR μV)/(r α γ) (*This is the
      condition that the veg eq is actually psositive*) && k2 < (r α γ - μR μV)/(r α γ)
     (*This is the condition that the veg2 eq is actually psositive*) &&
     θS (k1 (-r α γ + k1 r α γ + μR μV))/((-1 + k1) r μS) > μP (*The condition that the
     k1 strain can persist as a metapopulation on its own*)) /.
    {α → .5, μR → 0.1, μV → .2, μS → .05, θS → 0.35, γ → .5, μP → 0.5, r → 1},
   {k1, 0, 1}, {k2, 0, 1}, PlotPoints → 100,
   PlotStyle → Yellow], Plot[k1, {k1, 0, 1}]]
```

Out[ ]=

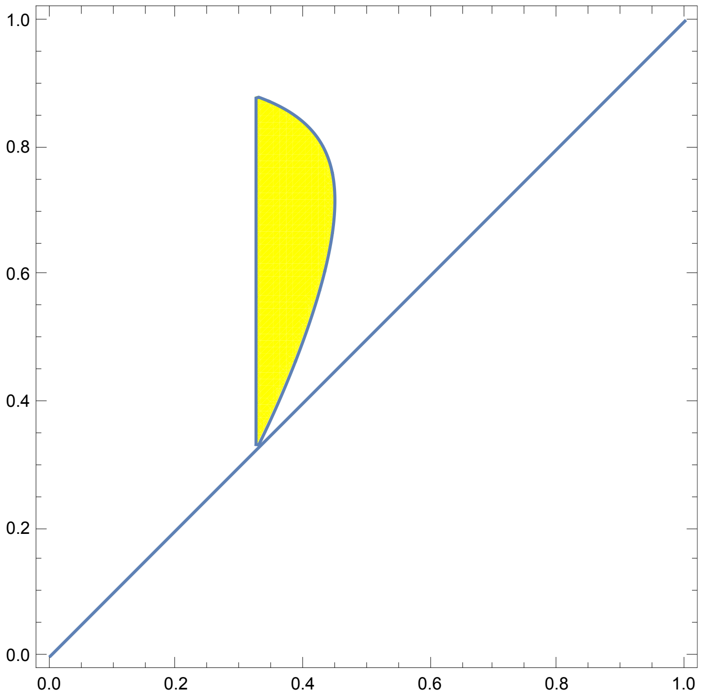

In[ ]:=
```
vegInvPlotCo = Show[RegionPlot[
  (k1 > k2 (*Just need to make sure that k2 is actually the better competitor*) &&
    k1 < (r α γ - μR μV)/(r α γ) (*This is the condition that the
     veg eq is actually psositive*) && k2 < (r α γ - μR μV)/(r α γ)
    (*This is the condition that the veg2 eq is actually psositive*) &&
    θS (k1 (-r α γ + k1 r α γ + μR μV))/((-1 + k1) r μS) > μP (*The condition that the
     k1 strain can persist as a metapopulation on its own*) &&
    θS (k2 (-r α γ + k2 r α γ + μR μV))/((-1 + k2) r μS) > μP (*The condition that the
     k2 strain can persist as a metapopulation on its own*) &&
    θS (k1 (-r α γ + k1 r α γ + μR μV))/((-1 + k1) r μS) > (θS (k2 (-r α γ + k2 r α γ + μR μV))/((-1 + k2) r μS))^2 / μP
    (*And they have coexistence*)
   ) /. {α → .5, μR → 0.1, μV → .2, μS → .05, θS → 0.35, γ → .5, μP → 0.5, r → 1},
  {k1, 0, 1}, {k2, 0, 1}, PlotPoints → 100, PlotStyle → Green], Plot[k1, {k1, 0, 1}]]
```

Out[ ]=

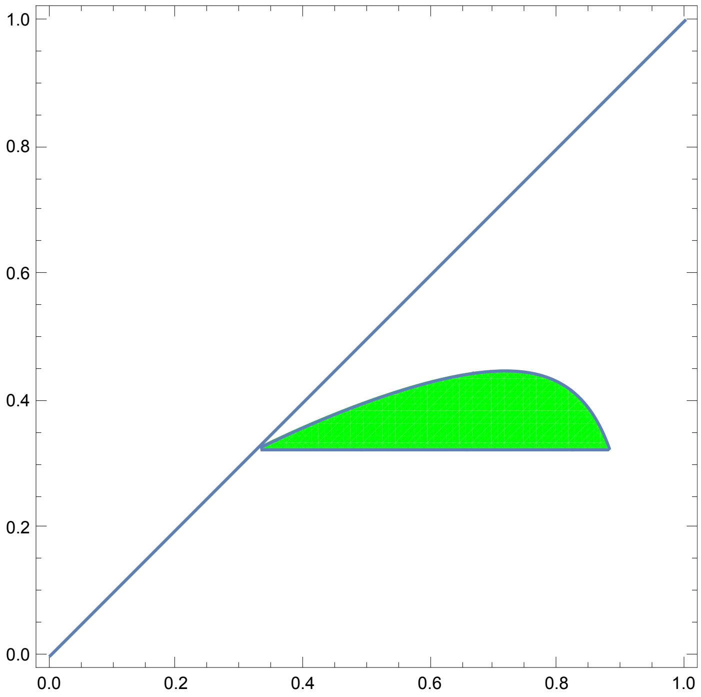

In[ ]:=

```
vegInvPlot = Show[RegionPlot[
  (k1 > k2 (*Just need to make sure that k2 is actually the better competitor*) &&
    k1 < (r α γ - μR μV)/(r α γ) (*This is the condition that the
     veg eq is actually psositive*) && k2 < (r α γ - μR μV)/(r α γ)
    (*This is the condition that the veg2 eq is actually psositive*) &&
    θS (k1 (-r α γ + k1 r α γ + μR μV))/((-1 + k1) r μS) > μP (*The condition that the
     k1 strain can persist as a metapopulation on its own*) &&
    θS (k2 (-r α γ + k2 r α γ + μR μV))/((-1 + k2) r μS) > μP (*The condition that the k2
     strain can persist as a metapopulation on its own*) &&
    θS (k1 (-r α γ + k1 r α γ + μR μV))/((-1 + k1) r μS) < (θS (k2 (-r α γ + k2 r α γ + μR μV))/((-1 + k2) r μS))^2 / μP
    (*And they have coexistence*)
   ) /. {α → .5, μR → 0.1, μV → .2, μS → .05, θS → 0.35, γ → .5, μP → 0.5, r → 1},
  {k1, 0, 1}, {k2, 0, 1}, PlotPoints → 100, PlotStyle → Red], Plot[k1, {k1, 0, 1}]]
```

Out[ ]=

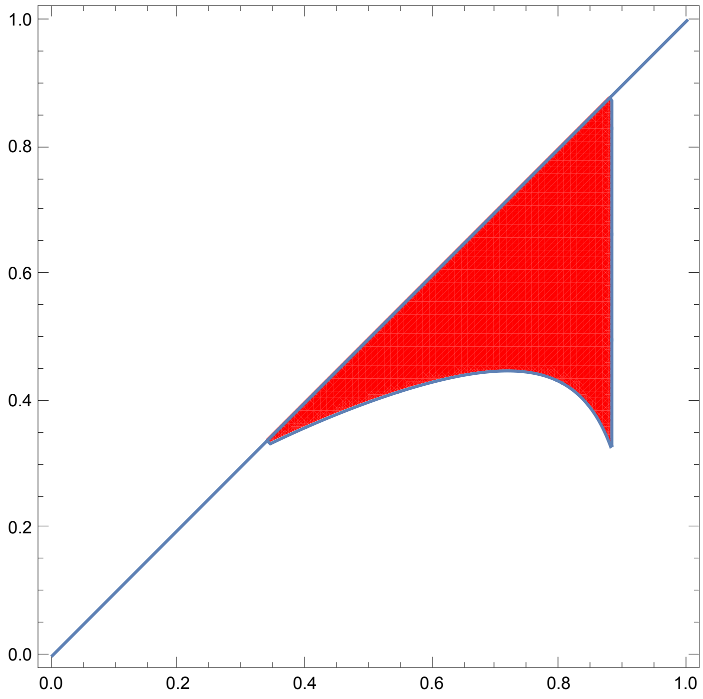

In[ ]:=
```
PIPPlot = Show[vegInvPlot, vegInvPlotCo, spoInvPlot,
  Framed → True, FrameLabel → {"Resident ρ̂", "Mutant ρ"}, BaseStyle →
   {FontSize → 12, FontWeight → Plain, FontFamily → "Helvetica"}, ImageSize → 300]
```

Out[ ]=

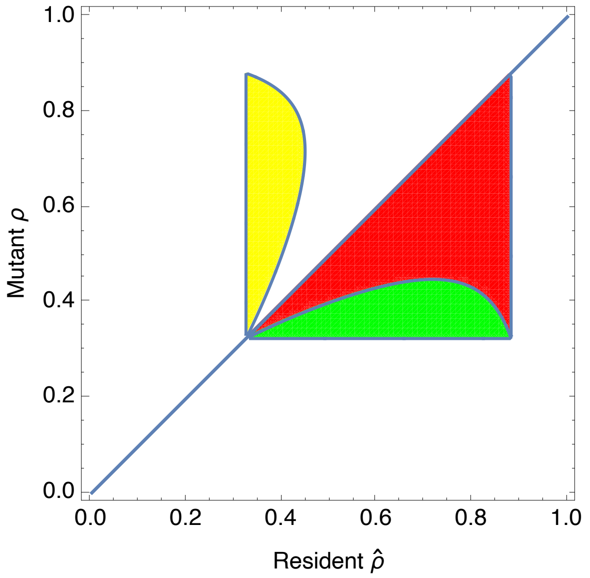


In[ ]:=

$\rho_{cH} = 1 - \sqrt{\frac{\mu R\, \mu V}{r\, \alpha\, \gamma}}$ /. {α → .5, μR → 0.1, μV → .2, γ → .5, r → 1};

$\rho_{cL}$ = 0.45;

In[ ]:=
```
lineStyle = {Thick, Purple, Dashed};
line1 = Line[{{ρcH, 0}, {ρcH, ρcH}}];
line2 = Line[{{0, ρcH}, {ρcH, ρcH}}];
line3 = Line[{{ρcL, 0}, {ρcL, ρcH}}];
line4 = Line[{{0, ρcL}, {ρcH, ρcL}}];

PIPPlot = Show[vegInvPlot, vegInvPlotCo, spoInvPlot,
  Framed → True, FrameLabel → {"Resident ρ̂", "Mutant ρ"},
  BaseStyle → {FontSize → 12, FontWeight → Plain, FontFamily → "Helvetica"},
  ImageSize → 300, Epilog → {Directive[lineStyle], line1, line2, line3, line4,
    Directive[Black], Text["ρcL", {ρcL + 0.05, 0.01}], Text["ρcH", {ρcH + 0.05, 0.01}],
    Text["ρcL", {0.05, ρcL + 0.03}], Text["ρcH", {0.05, ρcH + 0.03}]}]
```

Out[ ]=

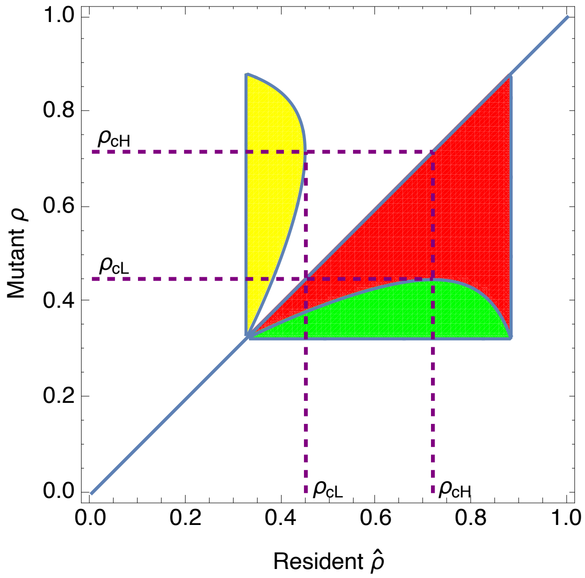


In[ ]:=
```
Export["PIPPlot.pdf", Show[PIPPlot,
  Prolog → {Opacity[0], Texture[{{{0, 0, 0, 0}}}], VertexTextureCoordinates →
     {{0, 0}, {1, 0}, {1, 1}}, Polygon[{{0, 0}, {.1, 0}, {.1, .1}}]
   }]]
```

Out[ ]=

```
PIPPlot.pdf
```

#### Dynamic PIP plots

In[ ]:=
```
Manipulate[Show[Plot[k1, {k1, 0, 1}],
  RegionPlot[(θS (k2 (-r α γ + k2 r α γ + μR μV))/((-1 + k2) r μS) > (θS (k1 (-r α γ + k1 r α γ + μR μV))/((-1 + k1) r μS))^2 / μP
     (*k2 has a sporulation advantage that is large enough to allow
```

```
      coexistence*) && k2 > k1 (*Just need to make sure that k1 is
                                                  r α γ - μR μV
      actually the better competitor*) && k1 < ───────────────── (*This is the
                                                    r α γ
                                                                      r α γ - μR μV
      condition that the veg eq is actually psositive*) && k2 < ─────────────────
                                                                         r α γ
    (*This is the condition that the veg2 eq is actually psositive*) &&
        k1 (-r α γ + k1 r α γ + μR μV)
    θS ──────────────────────────────── > μP (*The condition that the
              (-1 + k1) r μS

    k1 strain can persist as a metapopulation on its own*)) /.

  {α → .5, μR → 0.1, μV → .2, μS → .05, θS → 0.31, γ → .5, r → 1},
 {k1, 0, 1}, {k2, 0, 1}, PlotPoints → 100,
 PlotStyle → Yellow], RegionPlot[

 (k1 > k2 (*Just need to make sure that k2 is actually the
                                          r α γ - μR μV
      better competitor*) && k1 < ───────────────── (*This is the condition
                                             r α γ
                                                            r α γ - μR μV
      that the veg eq is actually psositive*) && k2 < ─────────────────
                                                               r α γ
    (*This is the condition that the veg2 eq is actually psositive*) &&
        k1 (-r α γ + k1 r α γ + μR μV)
    θS ──────────────────────────────── > μP (*The condition that the
              (-1 + k1) r μS
      k1 strain can persist as a metapopulation on its own*) &&
        k2 (-r α γ + k2 r α γ + μR μV)
    θS ──────────────────────────────── > μP (*The condition that the
              (-1 + k2) r μS
      k2 strain can persist as a metapopulation on its own*) &&
        k1 (-r α γ + k1 r α γ + μR μV)       (    k2 (-r α γ + k2 r α γ + μR μV) )
    θS ──────────────────────────────── >  ( θS ──────────────────────────────── ) ^2 / μP
              (-1 + k1) r μS                 (         (-1 + k2) r μS          )
    (*And they have coexistence*)

  ) /. {α → .5, μR → 0.1, μV → .2, μS → .05, θS → 0.31, γ → .5, r → 1},

 {k1, 0, 1}, {k2, 0, 1}, PlotPoints → 100, PlotStyle → Green], RegionPlot[

 (k1 > k2 (*Just need to make sure that k2 is actually the
                                          r α γ - μR μV
      better competitor*) && k1 < ───────────────── (*This is the condition
                                             r α γ
                                                            r α γ - μR μV
      that the veg eq is actually psositive*) && k2 < ─────────────────
                                                               r α γ
    (*This is the condition that the veg2 eq is actually psositive*) &&
        k1 (-r α γ + k1 r α γ + μR μV)
    θS ──────────────────────────────── > μP (*The condition that the
              (-1 + k1) r μS
```

```
      k1 strain can persist as a metapopulation on its own*) &&
     θS (k2 (-r α γ + k2 r α γ + μR μV)) / ((-1 + k2) r μS) > μP (*The condition that the k2
      strain can persist as a metapopulation on its own*) &&
     θS (k1 (-r α γ + k1 r α γ + μR μV)) / ((-1 + k1) r μS) < (θS (k2 (-r α γ + k2 r α γ + μR μV)) / ((-1 + k2) r μS))^2 / μP
     (*And they have coexistence*)
    ) /. {α → .5, μR → 0.1, μV → .2, μS → .05, θS → 0.31, γ → .5, r → 1},
  {k1, 0, 1}, {k2, 0, 1}, PlotPoints → 100, PlotStyle → Red]], {μP, 0.1, 0.9}]
```

#### Continuous ρ coexistence model

In[∘]:=
```
Clear[r, α, γ, μR, μV, μS, klow, khigh, eps, p, θS, ρmin, plow]
```

In[∘]:=
```
c[k1_] := θS (k1 (-r α γ + k1 r α γ + μR μV)) / ((-1 + k1) r μS)
```

Find the solution to the ODE, including a boundary condition for the

In[∘]:=
```
DSolve[
 {D[p[ρ], ρ] == p[ρ] (2 c[ρ] × D[c[ρ], {ρ, 2}] - 3 D[c[ρ], ρ]^2) / (2 c[ρ] × D[c[ρ], ρ]),
  p[ρmin] == plow}, p[ρ], ρ]
```

Out[∘]=

$$\left\{\left\{p[\rho] \to \frac{\text{plow}\left(r\,\alpha\,\gamma-\mu R\,\mu V-2\,r\,\alpha\,\gamma\,\rho+r\,\alpha\,\gamma\,\rho^{2}\right)\sqrt{1-\rho\text{min}}\;\rho\text{min}^{3/2}\left(-r\,\alpha\,\gamma+\mu R\,\mu V+r\,\alpha\,\gamma\,\rho\text{min}\right)^{3/2}}{\sqrt{1-\rho}\;\rho^{3/2}\left(-r\,\alpha\,\gamma+\mu R\,\mu V+r\,\alpha\,\gamma\,\rho\right)^{3/2}\left(r\,\alpha\,\gamma-\mu R\,\mu V-2\,r\,\alpha\,\gamma\,\rho\text{min}+r\,\alpha\,\gamma\,\rho\text{min}^{2}\right)}\right\}\right\}$$

We rewrite this in a way that is easier to read as:

In[∘]:=
```
pdisteq := plow (Sqrt[1 - ρmin] (ρmin)^(3/2) (r (1 - ρ)^2 α γ - μR μV) (-(1 - ρmin) r α γ + μR μV)^(3/2)) /
   (Sqrt[1 - ρ] ρ^(3/2) ((1 - ρmin)^2 r α γ - μR μV) (-r (1 - ρ) α γ + μR μV)^(3/2))
```

Solving for the portion that is dependent on ρ is equal to zero we get

In[∘]:=
```
Solve[(r α γ - 2 r ρ α γ + r ρ^2 α γ - μR μV) == 0, ρ] // Simplify
```

Out[∘]=

$$\left\{\left\{\rho \to 1-\frac{\sqrt{\mu R}\,\sqrt{\mu V}}{\sqrt{r}\,\sqrt{\alpha}\,\sqrt{\gamma}}\right\},\left\{\rho \to 1+\frac{\sqrt{\mu R}\,\sqrt{\mu V}}{\sqrt{r}\,\sqrt{\alpha}\,\sqrt{\gamma}}\right\}\right\}$$

Here we show that the extreme point in the yellow region is equal to xcrit. We first solve for the value of k1 that defines the edge of the yellow region, which gives us a relationship between resident and invader sporulation strategy. We then take the derivative of the critical value of k1 as a function of k2.

In[ ]:=
```
dtmp =
  ρ1 /. Solve[sc (ρ2 (-r α γ + ρ2 r α γ + μR μV))/((-1 + ρ2) r μS) == (sc (ρ1 (-r α γ + ρ1 r α γ + μR μV))/((-1 + ρ1) r μS))^2 / μP,
      ρ1][[3]] // D[#, ρ2] &;
```

We now substitute in our value of xcrit and simplify to show that the derivative is 0 at xcrit, implying that it is at the extremum of the yellow region.

In[ ]:=
```
Simplify[dtmp /. ρ2 → 1 - Sqrt[(μR μV)/(r α γ)]]
```

Out[ ]=
```
0
```

#### Plot the equilibrium density of the different strategies.

Find the minimum value of ρ that supports metapopulation persistence.

In[ ]:=
```
Solve[((ρ θS ((-1 + ρ) r α γ + μR μV))/((-1 + ρ) r μS) == μP) /.
   {α → .5, μR → 0.1, μV → .2, μS → .05, θS → 0.35, γ → .5, μP → 0.25, r → 1}, ρ]
```

Out[ ]=
```
{{ρ → 0.157852}, {ρ → 0.905005}}
```

Calculate the equilibrium density of the strain with the strategy of ρmin

In[ ]:=
```
ρmin = 0.18;
```

In[ ]:=
```
plow = pp /. Solve[-μP pp + pp c[ρmin] (1 - pp) == 0, pp][[2]] /.
   {α → .5, μR → 0.1, μV → .2, μS → .05, θS → 0.35, γ → .5, μP → 0.25, r → 1}
```

Out[ ]=
```
0.120549
```

In[ ]:=
```
Plot[
 pdisteq /. {α → .5, μR → 0.1, μV → .2, μS → .05, θS → 0.35, γ → .5, μP → 0.25, r → 1} //
   Re // If[# > 0, #, 0] &, {ρ, ρmin, 1}]
```

Out[ ]=

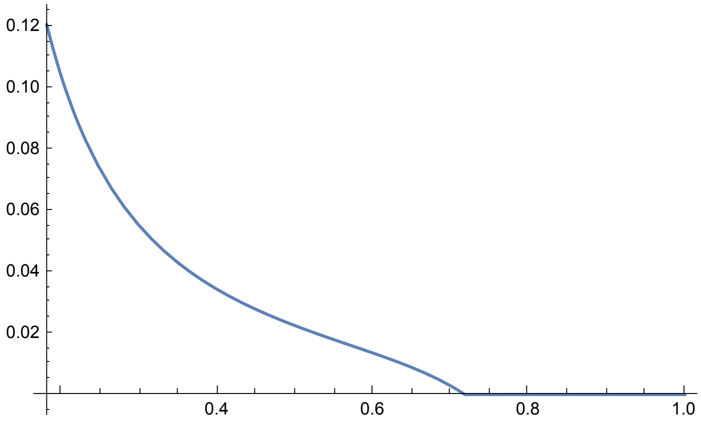


So I had it backwards, its the other shape that determines the max value. Interestingly, the maximum sporulation rate included does not depend on the patch extinction rate.

In[ ]:=
```
Manipulate[Plot[1 - Sqrt[x/y], {x, 0, 1}], {y, 0.1, 1}]
```

Out[ ]=

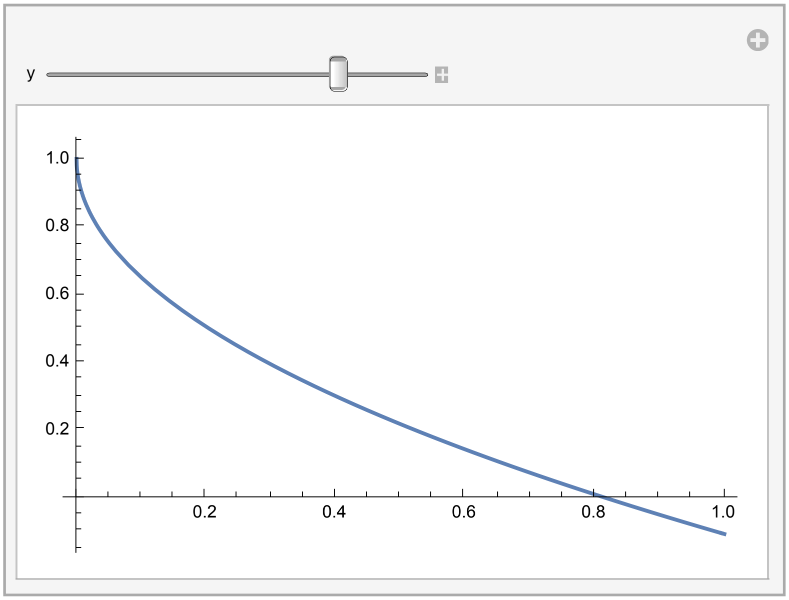

In[ ]:=
```
dplots = Table[
  ρmax = 1 - Sqrt[(μR μV)/(r α γ)] /. {α → .5, μR → 0.1, μV → .2, μS → .05, θS → 0.35, γ → .5, r → 1};
  xtmp = ρ /. (Solve[((ρ θS ((-1 + ρ) r α γ + μR μV))/((-1 + ρ) r μS) == μP) /. {α → .5, μR → 0.1,
        μV → .2, μS → .05, θS → 0.35, γ → .5, μP → i, r → 1}, ρ])[[1]];
  ρmin = (xtmp * 0.9 + ρmax * 0.1);
  plow = pp /. Solve[-μP pp + pp c[ρmin] (1 - pp) == 0, pp][[2]] /.
     {α → .5, μR → 0.1, μV → .2, μS → .05, θS → 0.35, γ → .5, μP → i, r → 1};
  Show[Plot[
    pdisteq /. {α → .5, μR → 0.1, μV → .2, μS → .05, θS → 0.35, γ → .5, μP → i, r → 1} //
      Re // If[# > 0, #, 0] &, {ρ, ρmin, 1}, PlotRange → {{0, 0.75}, {0, 1}}],
   Plot[0, {ρ, 0, ρmin / 1.1}, PlotStyle → {Thick, Red}]], {i, 0.01, 0.75, 0.01}];
```

In[ ]:=
```
ListAnimate[dplots]
```

Out[ ]=

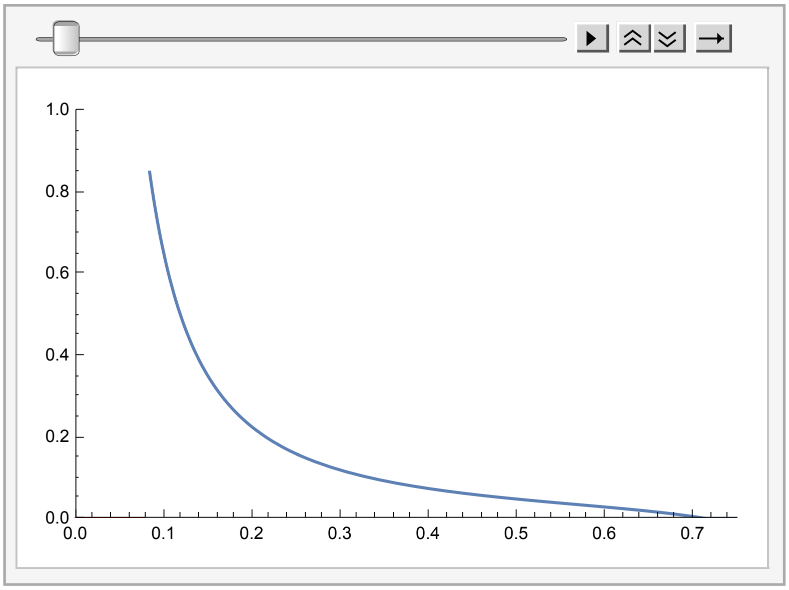


Create the functions for changing levels of $\mu P$

In[ ]:= 
```
dplots = Table[

    ρmax = 1 - Sqrt[(μR μV)/(r α γ)] /. {α → .5, μR → 0.1, μV → .2, μS → .05, θS → 0.35, γ → .5, r → 1};
    xtmp = ρ /. (Solve[((ρ θS ((-1 + ρ) r α γ + μR μV))/((-1 + ρ) r μS) == μP) /. {α → .5, μR → 0.1,
            μV → .2, μS → .05, θS → 0.35, γ → .5, μP → i, r → 1}, ρ]) [[1]];
    ρmin = (xtmp * 0.8 + ρmax * 0.2);
    plow = pp /. Solve[-μP pp + pp c[ρmin] (1 - pp) == 0, pp] [[2]] /.
       {α → .5, μR → 0.1, μV → .2, μS → .05, θS → 0.35, γ → .5, μP → i, r → 1};
    pdisteq /. {α → .5, μR → 0.1, μV → .2, μS → .05, θS → 0.35, γ → .5, μP → i, r → 1} //
       Re // If[(# > 0) && (ρ > ρmin), #, -1] &, {i, 0.05, 0.75, 0.1}];
```

In[ ]:= 
```
Table[i, {i, 0.05, 0.75, 0.1}]
```

Out[ ]=
```
{0.05, 0.15, 0.25, 0.35, 0.45, 0.55, 0.65, 0.75}
```

In[ ]:=
```
Olevel = 0.5;
Plot[{dplots[[1]], dplots[[4]], dplots[[7]]}, {ρ, 0, 1}, PlotRange → {0, 1} ,
 PlotStyle → {{Red, Thickness[0.01], Opacity[Olevel]}, {Purple, Thickness[0.01],
    Opacity[Olevel]}, {Green, Thickness[0.01], Opacity[Olevel]}},
 Frame → True, FrameLabel → {"ρ", "Density"}, BaseStyle →
  {FontSize → 12, FontWeight → Plain, FontFamily → "Helvetica"}, ImageSize → 300,
 Epilog → {Arrow[{{{0.325, 0.78}, {0.19, 0.66}}, {{0.54, 0.48}, {0.39, .23}},
     {{0.28, 0.07}, {0.52, 0.07}}}], Text["μP=0.05", {0.42, 0.8}],
   Text["μP=0.35", {0.64, 0.5}], Text["μP=0.65", {0.17, 0.07}]}]
```

Out[ ]=

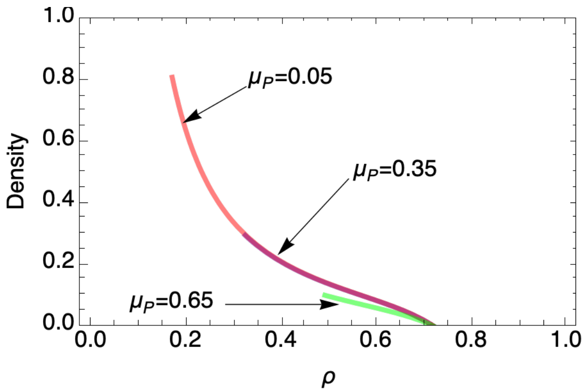


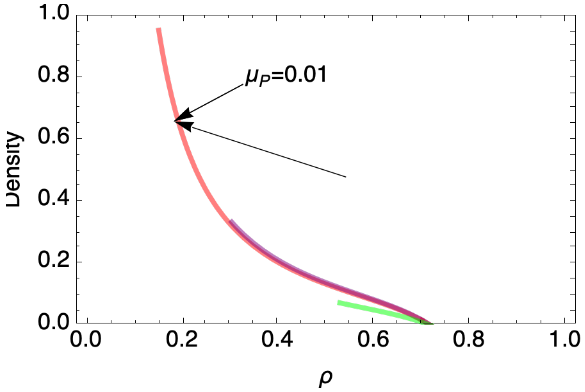

In[ ]:=
```
Show[dplots[[{1, 4, 8}]], PlotRange → {0, 1} ,
 PlotStyle → {Red, Blue, Green}, Frame → True, FrameLabel → {"ρ", "Density"}]
```

Out[ ]=

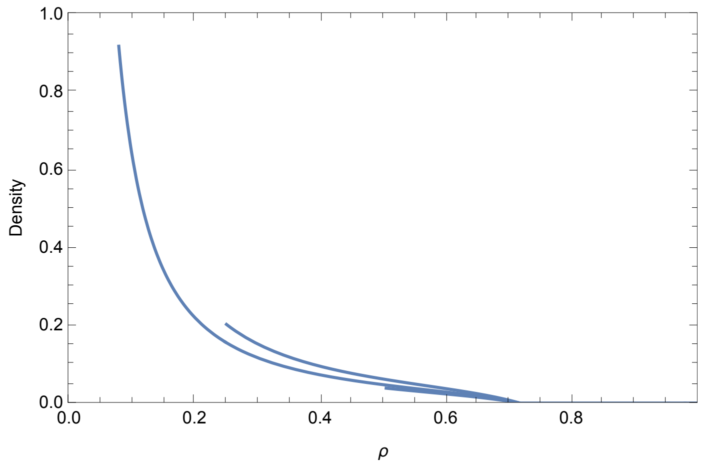


Calculate integral of patch frequency

In[ ]:=
```
dpatchOcc = Table[
  ρmax = 1 - Sqrt[(μR μV)/(r α γ)] /. {α → .5, μR → 0.1, μV → .2, μS → .05, θS → 0.35, γ → .5, r → 1};
  xtmp = ρ /. (Solve[((ρ θS ((-1 + ρ) r α γ + μR μV))/((-1 + ρ) r μS) == μP) /. {α → .5, μR → 0.1,
        μV → .2, μS → .05, θS → 0.35, γ → .5, μP → i, r → 1}, ρ])[[1]];
  ρmin = (xtmp * 0.9 + ρmax * 0.1);
  plow = pp /. Solve[-μP pp + pp c[ρmin] (1 - pp) == 0, pp][[2]] /.
    {α → .5, μR → 0.1, μV → .2, μS → .05, θS → 0.35, γ → .5, μP → i, r → 1};
  {i, NIntegrate[pdisteq /. {α → .5, μR → 0.1, μV → .2,
      μS → .05, θS → 0.35, γ → .5, μP → i, r → 1} // Re, {ρ, ρmin, ρmax}]}
  , {i, 0.1, 0.75, 0.01}];
```

In[ ]:= 
```
ListPlot[%]
```

Out[ ]=

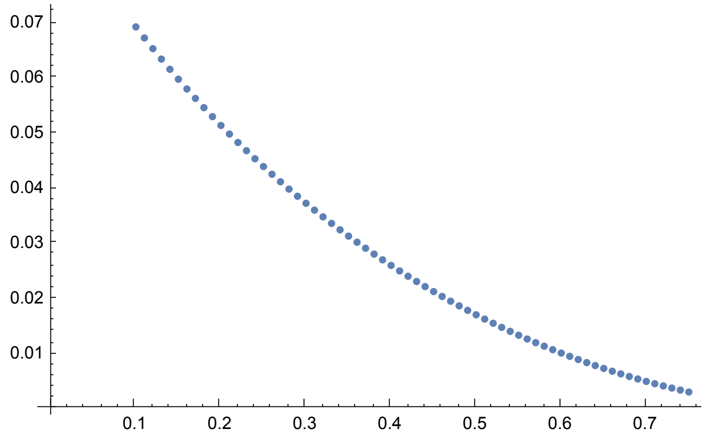
